# Deconvolute individual genomes from metagenome sequences through read clustering

**DOI:** 10.1101/620666

**Authors:** Kexue Li, Lili Wang, Lizhen Shi, Li Deng, Zhong Wang

## Abstract

**Motivation:** Metagenome assembly from short next-generation sequencing data is a challenging process due to its large scale and computational complexity. Clustering short reads before assembly offers a unique opportunity for parallel downstream assembly of genomes with individualized optimization. However, current read clustering methods suffer either false negative (under-clustering) or false positive (over-clustering) problems.

**Results:** Based on a previously developed scalable read clustering method on Apache Spark, SpaRC, that has very low false positives, here we extended its capability by adding a new method to further cluster small clusters. This method exploits statistics derived from multiple samples in a dataset to reduce the under-clustering problem. Using a synthetic dataset from mouse gut microbiomes we show that this method has the potential to cluster almost all of the reads from genomes with sufficient sequencing coverage. We also explored several clustering parameters that deferentially affect genomes with various sequencing coverage.

**Availability:** https://bitbucket.org/berkeleylab/jgi-sparc/.

## 1 INTRODUCTION

The shortgun sequencing of all DNA/RNA from clinical or environmental samples, or metagenomics sequencing, holds the key to comprehensively understand the structure, dynamics and interactions of underlying microbial communities and their implication to health and environment (Chiu and Miller, 2019; Tringe and Rubin, 2005; Thomas et al., 2012). As these samples often consist of thousands of different species with highly uneven richness, exceptional sequencing depth is required to study relatively rare species. As a result, except for a few cases (Brown et al., 2017), the majority of metagenome sequencing projects relied on cost-effective, short-read sequencing technologies like Illumina. These projects routinely produce a huge amount of data, ranging 100-1,000 giga-bases (Gb) or more (Howe et al., 2014; Shi et al., 2014). The largest project so far is the Tara Ocean Metagenomics project, where 7.2 tera-bases (Tb) was generated and the Prokaryote subset alone contains 28.8 billion short reads (Sunagawa et al., 2015).

As the majority of members of these microbial communities are unseen, assembling the short reads into draft genomes, or metagenome assembly, is a key step in metagenomics. Metagenome assembly pipelines have to deal with both the scale (billions of short, 100-250 bp reads) and the complexity problem (thousands of different species with a highly uneven abundance distribution). Most assemblers first assemble the short reads into longer contigs, then cluster the contigs into individual draft genomes through the binning process (Roumpeka et al., 2017; Kang et al., 2016). The assembly step in these software tools simultaneously tackles the computational and algorithmic challenges by constructing a huge *de bruijn* graph and subsequently partitions it in parallel (reviewed in Breitwieser et al. (2017); Quince et al. (2017)). These tools, including MegaHIT (Li et al., 2015), metaSpades (Nurk et al., 2017) and MetaHipmer (Georganas et al., 2018), have achieved considerable success and are widely used. To overcome the limitation of this “assembly-then-cluster” approach that does not allow optimization for individual genome assembly, a “cluster-then-assembly” alternative has recently been proposed. This strategy first clusters the reads based on their genome of origin, and then assemble each cluster individually (Guo et al., 2015; Shi et al., 2018). Some tools adopting such a strategy take advantage of the scalability and robustness of Apache ™ Hadoop (Guo et al., 2015) or Spark (Shi et al., 2018) platforms to construct and partition an overlap graph in parallel.

We previously reported that an Apache Spark™ based read clustering method, SpaRC, showed great potential in achieving good scalability and clustering performance (Shi et al., 2018). It can be flexibly deployed to different cloud or HPC computing environments. However, the demonstrated clustering success was limited to long read technologies. Even though SpaRC can form very pure clusters (low false positives), clustering short read datasets suffered a false negative problem, or one genome is clustered into many small clusters (under-clustering). This is not desirable as most of the metagenome datasets are based only on short read sequencing technologies. Clustering short reads to recover single genomes has been previously shown to be possible by a latent strain analysis approach (LSA, Cleary et al. (2015)). However, clustering metagenome reads directly based on k-mer statistics across multiple samples is very challenging because these statistics are very noisy and may not be very reliable.

In this paper, we describe a new method to target the under-clustering problem of SpaRC by exploiting statistics derived from multiple, independent samples in a short-read dataset. This method first robustly estimates the abundance of each cluster using a set of short, reliable k-mers, and then calculates the similarity among the clusters and uses it to construct a graph of clusters. Finally, it partitions the cluster graph to obtain larger clusters. We refer the new clustering algorithm developed here as “global clustering”, as it deals with cross-sample information from the entire dataset. Conversely the clustering algorithm in SpaRC reported previously is referred as “local clustering”, as it only deals with read overlap information. We implemented the global clustering algorithm on the Apache Spark platform to achieve data scalability and computing robustness. In addition, we adopted minimizers (Roberts et al., 2004) as a replacement for k-mers to improve computing and memory efficiency. We compared the clustering performance of the global clustering algorithm to the local clustering in the original SpaRC, using a synthetic mouse gut microbiome dataset from the CAMI2 project (Sczyrba et al., 2017). Several clustering parameters were also explored to gauge their influence on global clustering performance.

## MATERIALS AND METHODS

### Algorithm overview

An overview of the entire software is shown in Figure 1. The local clustering algorithm in the original SpaRC has been described in our previous work (Shi et al., 2018), below we only describe the improvements we made in this study.

**Figure 1.**
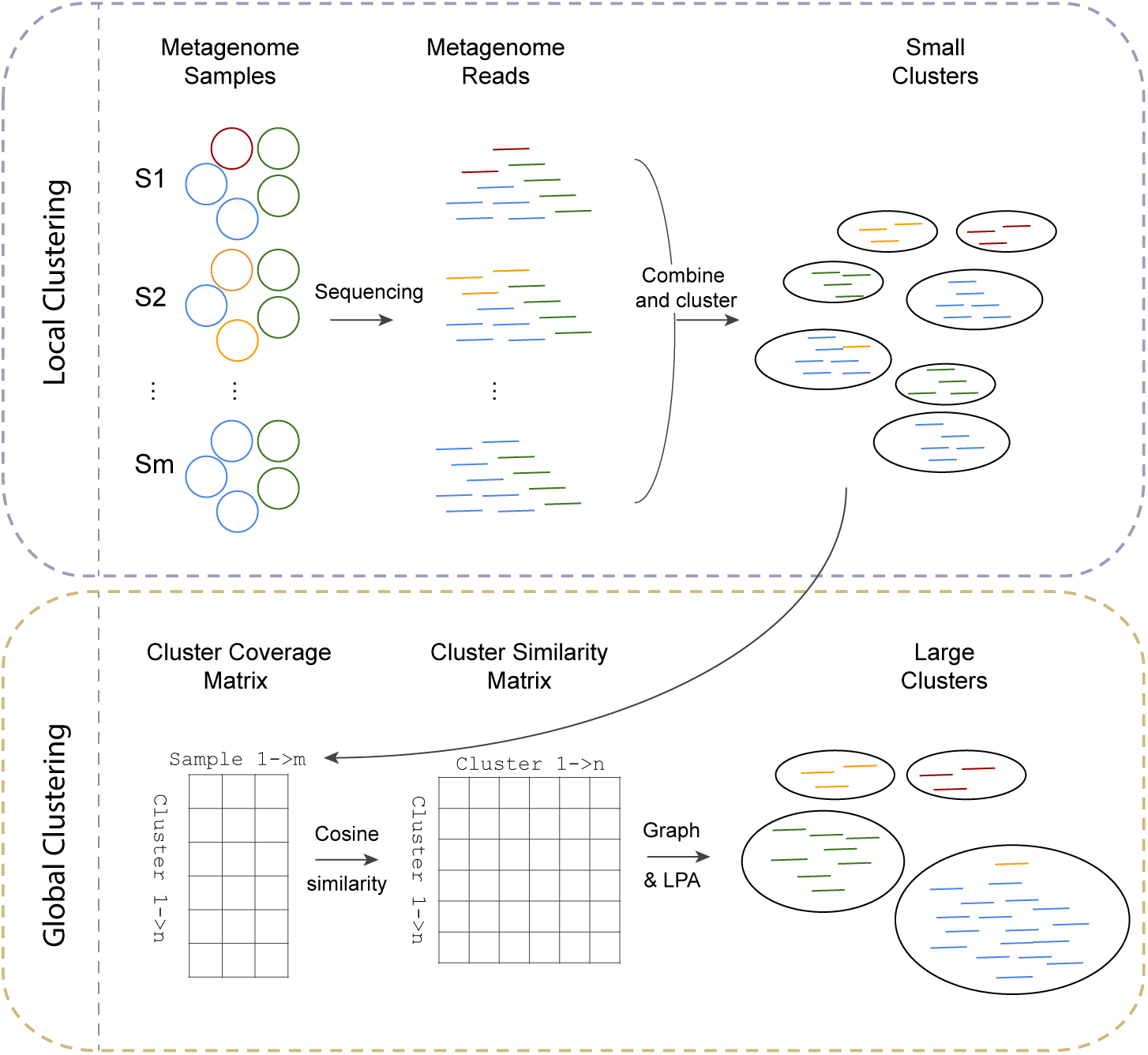
An overview of the entire software. (**Top**) The algorithm for local clustering: short reads sequences from multiple samples of a microbial communities (such as derived from different sample sites or times, S1, S2 … Sm) are combined and clustered using the scalable overlap-based clustering algorithm in SpaRC. Many small clusters are formed and reads from the same genomes scatter across many clusters (under-clustering). (**Bottom**) The algorithm for global clustering: First, sequencing coverage of each small cluster from the local clustering step is estimated and a cluster coverage matrix is derived. Second, a square similarity matrix is obtained by computing pair-wise cosine similarities between all clusters. Finally, a graph is constructed using clusters as nodes and their similarity as weighted edges. Larger clusters containing all the reads from individual genomes can be obtained by partitioning the graph using the Label Propagation Algorithm (LPA).

### Local clustering with minimizers

In SpaRC, the number of shared k-mers is used to estimate similarity between reads (Shi et al., 2018). As it takes 100-200x times more space after reads are transformed into k-mers and edges, SpaRC is neither space or time-efficient. To improve the computing efficiency, we implemented a new function to use minimizers (Roberts et al., 2004) instead of k-mers to estimate similarity between reads. As many adjacent shared k-mers can be represented by a single minimizer without losing sensitivity, in theory, the minimizer-based method should greatly reduce the memory requirement in SpaRC (as fewer k-mers and edges will be produced). In practice we did observe a 3.2-fold memory usage reduction, and 3.3-fold of speed-up (data not shown). It is worth noting that minimizers may not be applicable to uncorrected long reads from PacBio and Nanopore sequencing technologies due to high error rate.

### Global clustering

#### Estimating cluster coverage

Like the k-mer analysis done on the whole genome level (Chor et al., 2009; Lo and Chain, 2014), the k-mer frequency distribution calculated from a cluster should have a similar spectrum, where distinct k-mers form a main peak corresponding to sequencing coverage of the underlying genomic region, and erroneous k-mers pileup on the far left of the X-axis. For the pure clusters formed by SpaRC, they are likely to have a uni-modal k-mer spectrum, where the main peak after the minimum inflection point represents the sequencing coverage of the genome region where these reads are derived. We therefore select several k-mers around this peak, and use the median of their counts among each sample to estimate the coverage of this cluster in each sample. If two clusters belong to the same genome, then we expect their coverage should both be similar to the genome coverage. We term these k-mers as representative k-mers for each cluster. Because very small clusters do not generate reliable k-mer spectra, we only select clusters with more than 50 reads for further clustering. To avoid counting k-mers twice, we modified the original KMR function in SpaRC to track k-mer counts of each sample.

The cluster coverage information in stored in a *m* by *n* coverage matrix, where *m* is the number of samples and *n* is the number of read clusters.

#### Calculating similarity between clusters

In the above cluster coverage matrix, every cluster is represented by a vector of counts. If two clusters are derived from the same genome, we expect their vectors should be very similar. We chose cosine similarity to measure the similarity of cluster vectors as it is most commonly used in high-dimensional positive spaces. After all pairwise similarities are calculated, the coverage matrix is transformed into a *n* by *n* similarity matrix, where *n* is the number of clusters. We only keep the similarity exceeding a predefined threshold because of the sparsity nature of this matrix. This threshold parameter has a direct impact on clustering performance, as higher thresholds produce smaller but purer clusters. Optimal threshold can be learned by using labelled datasets.

#### Graph construction and partitioning

By using the cosine similarity calculated above as weighted edges and the clusters as nodes, a weighted graph can be constructed. This cluster graph can then be partitioned into big clusters the same way in the local clustering step by using the Label Propagation Algorithm(Raghavan et al., 2007).

### Datasets

#### The MBARC-26 microbial community

This mock dataset is a synthetic community with real-world sequence data(Singer et al., 2016). It contains Illumina reads from 23 bacteria and 3 Archaea species with known reference genomes. The sequence length is (90–150) *2 bp totaling 3.3 Gb (Table 1). This dataset was used as a toy dataset for testing local clustering with minimizers.

**Table 1.** Datasets used in this study.

| Dataset | CAMI2 | MBARC-26 |
| --- | --- | --- |
| # samples | 64 | 1 |
| # genomes | 791 | 26 |
| Read length(bp) | 2x150 | (90-150)x2 |
| Total size(Gb) | 320 | 3.3 |

#### CAMI2 mouse gut metagenome dataset

All the benchmark experiments on global clustering were done on a simulated dataset from the second CAMI Challenge (https://openstack.cebitec.uni-bielefeld.de:8080/swift/v1/CAMISIM_MOUSEGUT/). This dataset was simulated using 800 reference genomes derived from mouse gut microbiome, and it contains 64 samples with various genome coverage. Some relevant statistics of this dataset is shown in Table 1. A complete list of the organisms in this dataset is available at this URL (https://openstack.cebitec.uni-bielefeld.de:8080/swift/v1/CAMI_DATABASES/taxdump_cami2_toy.tar.gz).

### Computing environments

Read clustering experiments were performed on Amazon Web Service (AWS)’s Elastic MapReduce (EMR, emr-5.9.0). Depending on the size of the dataset, a number of r3.2xlarge instances were used to form a cluster, and the configuration details are shown in Table 2.

**Table 2.**
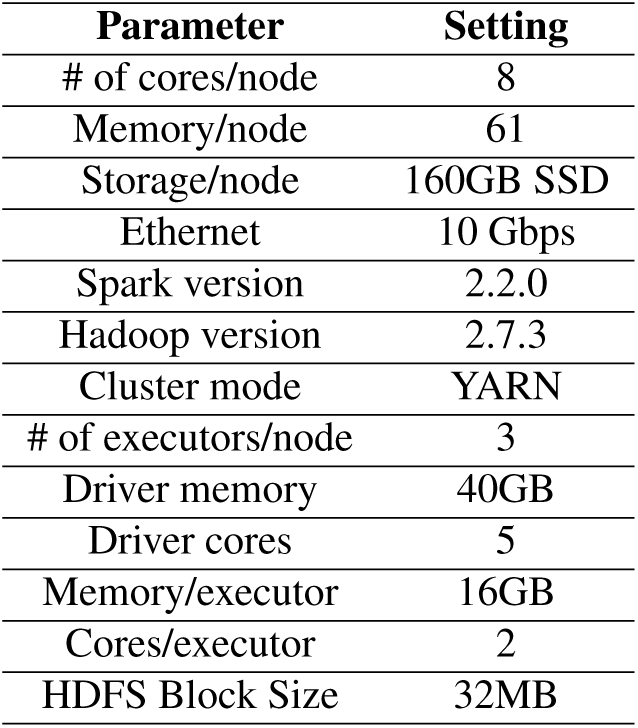
Configuration of AWS EMR.

| Parameter | Setting |
| --- | --- |
| # of cores/node | 8 |
| Memory/node | 61 |
| Storage/node | 160GB SSD |
| Ethernet | 10 Gbps |
| Spark version | 2.2.0 |
| Hadoop version | 2.7.3 |
| Cluster mode | YARN |
| # of executors/node | 3 |
| Driver memory | 40GB |
| Driver cores | 5 |
| Memory/executor | 16GB |
| Cores/executor | 2 |
| HDFS Block Size | 32MB |

## RESULTS

### Short read clustering performance is greatly improved with multiple samples from the same microbial community

In order to test whether multiple samples derived from the same microbial community could be leveraged to improve short read clustering performance, we designed a control dataset by taking 10% of the reads from 50 samples from the CAMI2 synthetic metagenome dataset (Materials and Methods). We clustered the reads using two clustering methods: in the “local clustering” method, we combined all the reads from the 50 samples for clustering and selected clusters with 100 reads or larger;in the “global clustering” method, we further applied the global clustering module to these clusters to form big clusters (Material and Methods). This labeled synthetic dataset enabled us to systematically compare the clustering performance between these two methods for cluster size, purity, completeness. The results are shown in Figure 2. We used the same purity and completeness metrics as in (Shi et al., 2018). Briefly, the purity of a cluster is defined as the percentage of reads from the dominant genome within the cluster, while the completeness of a cluster is defined as the percentage of all the reads from the dominant genome that are captured by the cluster. Because almost identical strains from the sample species were engineered in the dataset, both of the two metrics, especially purity, likely underestimate species-level clustering performance. For example, if a species has two closely related strains with equal number of reads, clusters derived from this species will have a purity of 50%. In this experiment, the default parameter set was used (specifically, k=31, m=18, min_shared_kmers=2, max_degree=25, representative k-mer count=9, and cosine threshold=0.925).

**Figure 2.**
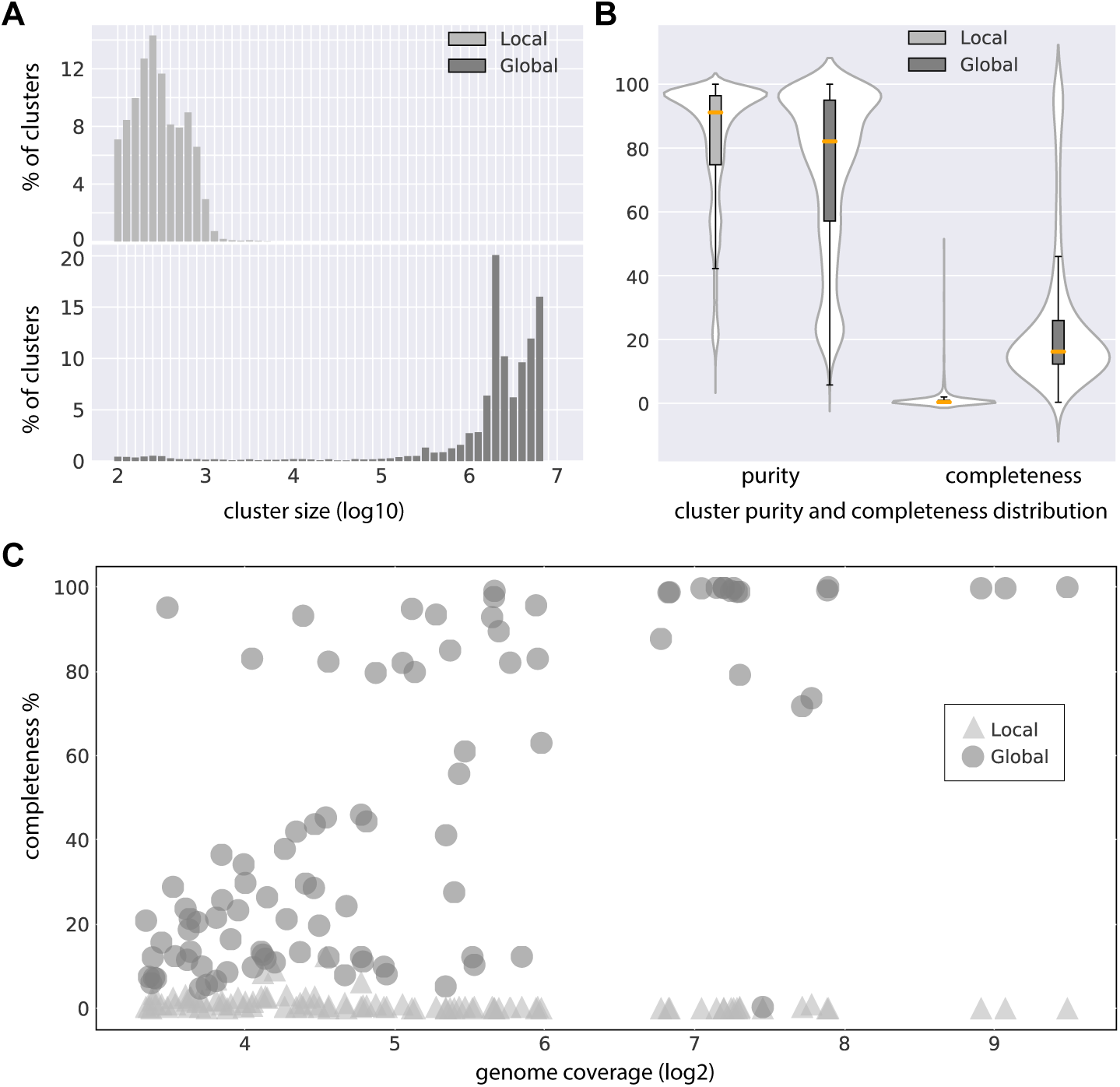
A comparison of the clustering performance between local clustering and global clustering. **A**. The distribution of clustered reads among the clusters for local clustering (top) and global clustering (bottom). x-axis is cluster size (log10) and y-axis is the percent of reads that are clustered at a given cluster size. **B**. Violin plots of cluster purity (left) and completeness (right). Local clustering metrics are in light gray, while global clustering metrics are in dark gray. **C**. A scatter plot of sequencing coverage of the genomes and their completeness from local clustering (light gray triangles) and global clustering (dark gray circles). x-axis is the sequencing coverage (log2) and y-axis is the completeness.

The local clustering step resulted in 74.92% of the reads clustered into many small clusters (n=233,768), with the largest cluster having only 19,473 reads. The majority of the clustered reads 95.81% are distributed in clusters with 1,000 reads or less (Figure 2A). In contrast, after the global clustering step the number of clusters is significantly reduced (n=9,324), 80.72% of reads are in clusters with 1,000,000 or more reads, and the largest cluster contains 9,351,200 reads. Consequently, the median completeness of the clusters from the global clustering is also 48.94 times larger than that from the local clustering (16.15% vs 0.33%, Figure 2B). The increase in completeness came without a big drop in purity (median purity from 91.15% to 82.07%), as most of the clusters are still pure (Figure 2B).

As the success of clustering of genome should heavily depend on its sequencing coverage, we next explored the relationship of cluster completeness as a function of genome sequencing coverage (Figure 2C). For local clustering, higher sequencing coverage seems to have little effect on cluster completeness. In contrast, higher sequencing coverage does translate into higher completeness, suggesting global clustering effectively leverages multiple sample statistics for read clustering. After the sequencing coverage reaches a sufficient threshold (100x, Figure 2C), the completeness of most genomes exceeds 80%.

### The performance of short read clustering can be improved by increasing the number of samples

We next explored the relationship between the number of samples in a dataset and the global clustering performance. Intuitively, more samples should enable more robust estimation of the similarity between clusters and lead to better clustering performance. By limiting the total size of the datasets to 25 Gb, we made several datasets with various number of samples (5,10,20,50) from the CAMI2 synthetic metagenome dataset (Materials and Methods). We obtained clusters from these datasets by running SpaRC with the same parameters (k=41, m=22, min_shared_kmers=2, min_read_per_cluster=50) for local clustering, and representative_kmer_count=100 for global clustering.

As in the previous section, we evaluated the purity and completeness of the resulting clusters. As shown in Figure 3, the median purity of the clusters slightly dropped as the number of sample increases, from the highest 96.96% (at n=5) to the lowest 88.89% (at n=20, Figure 3A). This is likely caused by the fact that more samples may contain more strain variation, and currently SpaRC can not distinguish very similar strains. In contrast, the completeness continuously increases with increasing number of samples (median completeness rises from 7.69% at n=5 to 18.75% at n=50, Figure 3B). This result supports the hypothesis that more samples enhance global clustering performance, likely due to better estimation of cluster similarity.

**Figure 3.**
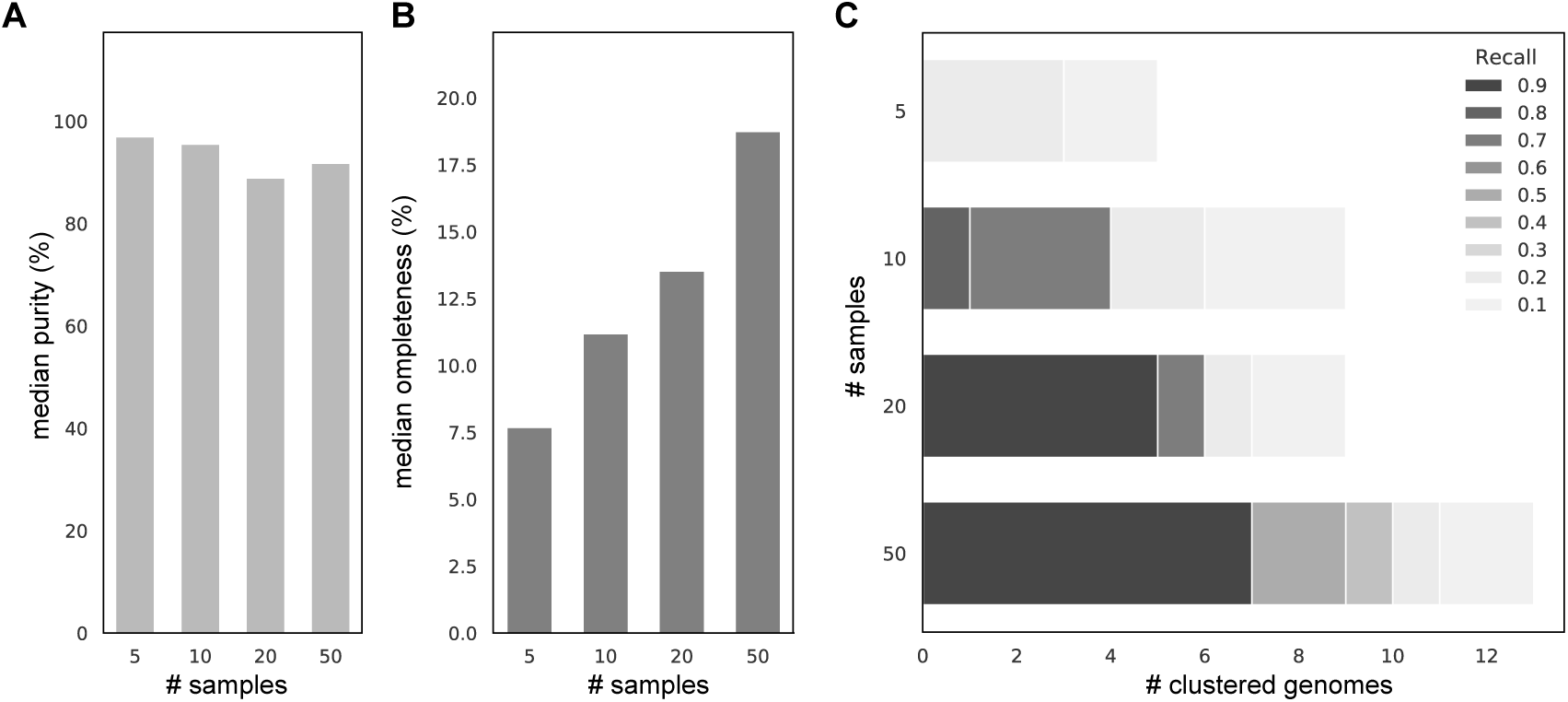
Clustering performance with different number of samples. **A**. Median purity comparison among different number of samples. x-axis is the number of samples and y-axis is median purity(light gray). **B**. Median completeness comparison among different number of samples. x-axis is the number of samples and y-axis is median completeness(dark gray). **C**. The number of clustered individual genomes with purity > 95% and completeness > 80% among different number of samples. Different color depth stands for different completeness. x-axis is the number of genomes and y-axis is the number of samples.

The ultimate goal of read clustering is to recover the complete set of any genome without any contamination from other genomes. In order to measure how many genomes can be recovered by read clustering, here we define “a clustered genome” as a read cluster that simultaneously satisfies two criteria: purity > 95% and completeness > 80%. It is worth noting that these criteria are very strict. As previously mentioned, strain-level heterogeneity can greatly reduce purity, and one small region larger than a read with no sequencing coverage, either due to statistical sampling or systematic sequencing biases, will greatly decrease completeness.

The clustered genomes from 5, 10, 20 an 50 samples are 0, 1, 5 and 7, respectively (Figure 3C). Since the ability to obtain a clustered genome depends on the sequencing coverage, and there are 19 genomes with at least 100X coverage, this translates to a recovery rate of 35% with 50 samples. Other genomes with lower sequencing coverage also benefit from more samples included in clustering, as Figure 3C shows the extent of recovery for all genomes with a sequence coverage > 10*X*. The detail of these 7 recovered genome can be found in the Supplemental Table 1.

### Parameters that may impact clustering performance

There are 10 parameters in the local clustering algorithm (Shi et al., 2018). In the global clustering step two more parameters are added, number of representative k-mers are used to estimate cluster abundance (*rp*) and cosine similarity threshold (*cs*). While some of these parameters only affect computing efficiency, there are four parameters that in theory may affect clustering accuracy: k-mer(*k*)/minimizer(*m*) length, min_shared_kmers among reads (*minsk*), *rp* and *cs*. In theory higher *k, minsk*, and *cs* all lead to smaller clusters with lower completeness but higher purity, and *vice versa*. Higher *rp* should make the estimation of cluster abundance more accurate with a small cost in computing efficiency. To explore the effect of these parameters on clustering accuracy, we ran SpaRC with different sets of parameters on the 25 Gb dataset with 50 samples. In each of the parameter set, only one parameter varies while the other three were held constant. The rest of parameters were used as their default values. We used several metrics including number of reads clustered (bigger is better), number of clusters formed (bigger is worse), median cluster purity, median cluster completeness and number of clustered genomes to measure clustering performance. The results are shown in Table 3.

**Table 3.**
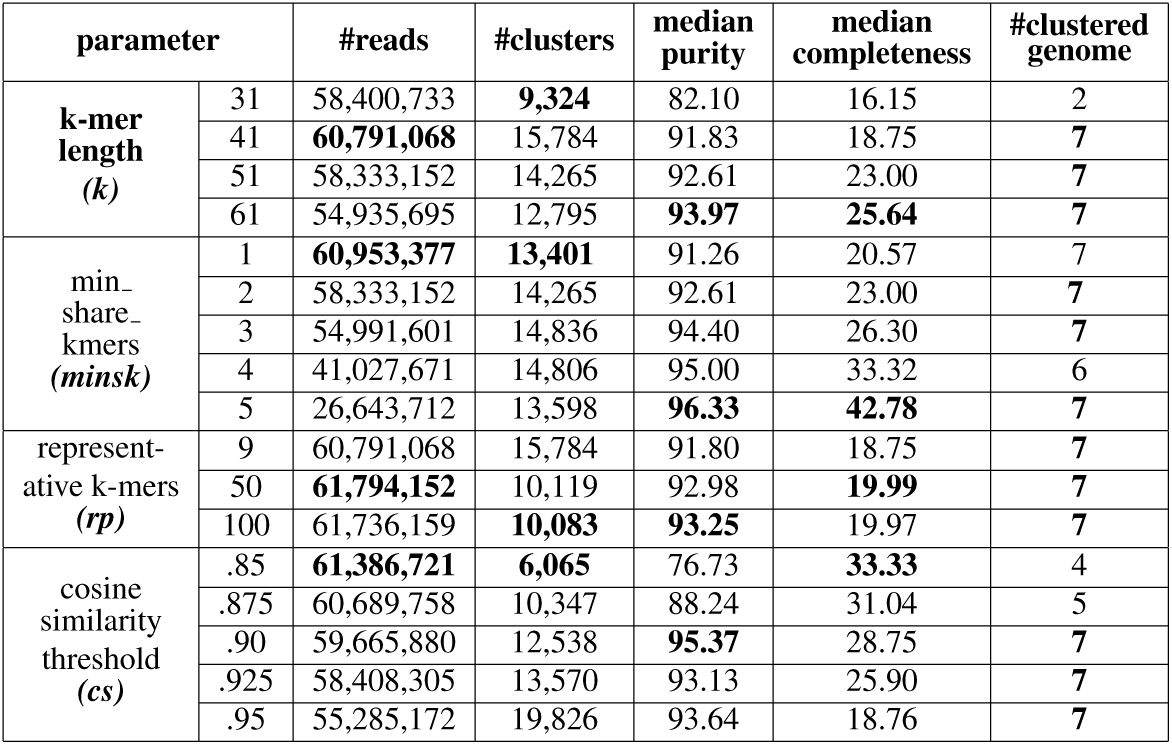
Clustering performance vs different parameters

Overall, there is no single best parameter that can maximize all metrics. In general, the most abundant genomes (number of clustered genomes) are less likely to be affected by these parameters, except a few extreme cases (k-mer is too small or cluster similarity threshold is too low). In those extreme cases over-clustering happened, as many fewer clusters formed with very low median purity. These parameters, except number of representative k-mers, can greatly affect the median purity and completeness. Longer k-mers and requiring more shared k-mers among reads increases median purity and completeness. These improvements are likely driven by better clusters from the genomes with medium to high sequencing coverage. If these parameters became very large, the number of reads clustered dramatically decreases (under-clustering), but these genomes do not seem to be affected. For example, the number of clustered reads drops from 60,953,377 at minsk=1 to only 26,643,712 at minsk=5, but the median completeness and purity reaches their peak. The un-clustered reads at high minsk presumably are derived from a lot of genomes with low sequencing coverage.

The number of representative k-mers used for cluster abundance estimation seems to have a very minor effect on clustering performance. If the number is too low, then the performance is slightly lower.

As expected, very low cluster similarity thresholds cause over-clustering, and very high ones lead to under-clustering. Same as the other parameters, the best parameter choice should be determined by the underlying scientific requirements to balance sensitivity and specificity.

## CONCLUSION AND DISCUSSION

In summary, we extended our previous work on the Apache Spark-based read clustering by exploiting species co-variation across different metagenome samples to improve clustering completeness. Using a complex control dataset with many samples, we showed the global clustering algorithm can dramatically improve both the cluster size and genome completeness with only short reads. Furthermore, this approach can potentially recover almost the full set of reads of microbial genomes with sufficient representation in the data, an ability influenced by several parameters including the number of available samples.

The read clustering problem is similar to the metagenome binning problem (Kang et al., 2015). Here we applied the abundance co-variation metric to improve clustering. The application of the tera-nucleotide-frequency (TNF) metric, however, is not straightforward, as TNF may not be reliably estimated from unassembled reads. Future analyses will be needed to integrate more information such as TNF into the clustering framework to reduce the requirement of many samples, given the fact that most of the metagenome shortgun sequencing experiments were carried out on single samples. As long read technologies (PacBio and Nanopore) are increasingly applied to metagenome sequencing, a hybrid clustering approach leveraging long reads may also greatly increase clustering performance. Long reads should be very helpful to increase clustering completeness as they span the regions where short reads have low coverage, and improve clustering purity strain as they can distinguish repeats and closely related species or strains.

The global clustering step added more parameters to the entire clustering pipeline. We have shown that clustering accuracy can be influenced by some of these parameters, including k-mer/minimizer size, minimum shared k-mers to detect overlap, and abundance similarity thresholds among clusters to construct the cluster graph. Deriving an optimal set of parameters is challenging because it is likely dependent on the underlying data characteristics and the scoring metrics. Further, searching for an optimal parameter set from a large parameter space on a large dataset is computational prohibitive. There isn’t an evident law to select the default parameters. This problem may be a good candidate for Bayesian optimization (Snoek et al., 2012; Hutter et al., 2011).

## ACKNOWLEDGMENT

We thank Alex Copeland for critical reading of the manuscript. The work was conducted by the US Department of Energy Joint Genome Institute. ZW’s work was supported by the U.S. Department of Energy, Office of Science, Office of Biological and Environmental Research under Contract No. DE-AC02-05CH11231.

