## Supplemental_Table1 for "Deconvolute individual genomes from metagenome sequences through read clustering"

| Table S1: Genome clusters with >95% purity vs the number of samples | | | | | | | | |
|---|---|---|---|---|---|---|---|---|
| #samples | ref_ID | ref_length(bp) | #ref_reads | ref_coverage | cluster_completeness | #cluster_ref_reads | #cluster_total_reads | cluster_purity |
| 50 | 4306262 | 2858672 | 2265169 | 237.71552 | **0.999872** | 2261914 | 2303113 | 0.982112 |
| 50 | 192222 | 3553509 | 1736429 | 146.595576 | **0.999809** | 1730860 | 1746950 | 0.99079 |
| 50 | 851733 | 2723877 | 6516886 | 717.751132 | **0.999091** | 6506143 | 6602789 | 0.985363 |
| 50 | 1684221 | 4587353 | 2166476 | 141.681445 | **0.999017** | 2154836 | 2191755 | 0.983156 |
| 50 | 318732 | 6627518 | 3387718 | 153.347814 | **0.997985** | 3372221 | 3496206 | 0.964537 |
| 50 | 999989 | 2685334 | 2116550 | 236.45662 | **0.995212** | 2099531 | 2191095 | 0.958211 |
| 50 | 4469576 | 3540263 | 1788460 | 151.553147 | **0.993745** | 1768773 | 1846979 | 0.957657 |
| 50 | 1108599 | 4628540 | 782815 | 50.738354 | 0.530813 | 384441 | 401749 | 0.956918 |
| 50 | 835900 | 3753503 | 682637 | 54.559994 | 0.502199 | 306589 | 313317 | 0.978527 |
| 50 | 187223 | 4326619 | 478367 | 33.16911 | 0.480858 | 148991 | 153929 | 0.96792 |
| 50 | 4394913 | 3715671 | 227396 | 18.359753 | 0.255171 | 5626 | 5768 | 0.975381 |
| 50 | 1107303 | 3565661 | 500706 | 42.127336 | 0.183182 | 71172 | 72356 | 0.983636 |
| 50 | 245324.1 | 3009832 | 463684 | 46.216932 | 0.122779 | 43894 | 45237 | 0.970312 |
| 20 | 318732 | 6627518 | 6055271 | 274.096773 | **0.99799** | 6043101 | 6185355 | 0.977001 |
| 20 | 999989 | 2685334 | 2339297 | 261.341457 | **0.986922** | 2308703 | 2395549 | 0.963747 |
| 20 | 4469576 | 3540263 | 3403100 | 288.37688 | **0.985376** | 3353332 | 3464161 | 0.968007 |
| 20 | 4306262 | 2858672 | 4610089 | 483.800415 | **0.974864** | 4494210 | 4589427 | 0.979253 |
| 20 | 4449524 | 3322005 | 1272011 | 114.87138 | **0.932702** | 1186407 | 1235294 | 0.960425 |
| 20 | 851733 | 2723877 | 3579674 | 394.255027 | 0.719848 | 2576821 | 2588950 | 0.995315 |
| 20 | 1684221 | 4587353 | 652380 | 42.663819 | 0.203349 | 132661 | 133544 | 0.993388 |
| 20 | 190114 | 3247548 | 859469 | 79.395501 | 0.180004 | 154708 | 156414 | 0.989093 |
| 20 | 4480176 | 3050713 | 371886 | 36.570402 | 0.158667 | 59006 | 60344 | 0.977827 |
| 10 | 4469576 | 3540263 | 4059551 | 344.004188 | **0.835594** | 3392135 | 3425222 | 0.99034 |
| 10 | 318732 | 6627518 | 8727296 | 395.048161 | 0.790555 | 6899407 | 7034156 | 0.980844 |
| 10 | 4306262 | 2858672 | 7367602 | 773.184402 | 0.729387 | 5373833 | 5655207 | 0.950245 |
| 10 | 1684221 | 4587353 | 1124793 | 73.558303 | 0.231096 | 259935 | 261074 | 0.995637 |
| 10 | 4424327 | 4400443 | 1279318 | 87.217446 | 0.217894 | 278756 | 283352 | 0.98378 |
| 10 | 851733 | 2723877 | 6108452 | 672.767383 | 0.198397 | 1211899 | 1217465 | 0.995428 |
| 10 | 4378740 | 3134201 | 2445375 | 234.066832 | 0.18916 | 462568 | 476798 | 0.970155 |
| 10 | 245324.1 | 3009832 | 183930 | 18.332917 | 0.14785 | 27194 | 27477 | 0.9897 |
| 10 | 835900 | 3753503 | 392123 | 31.340564 | 0.100892 | 39562 | 40437 | 0.978361 |
| 5 | 303652 | 1989200 | 343616 | 51.82224 | 0.282269 | 96992 | 101728 | 0.953444 |
| 5 | 999989 | 2685334 | 9233002 | 1031.492023 | 0.228715 | 2111725 | 2182437 | 0.9676 |
| 5 | 318732 | 6627518 | 16340903 | 739.684283 | 0.204774 | 3346189 | 3396696 | 0.985131 |
| 5 | denovo4581.0 | 2879979 | 369569 | 38.497052 | 0.193883 | 71653 | 73846 | 0.970303 |
| 5 | 1684221 | 4587353 | 2167300 | 141.735332 | 0.1193 | 258558 | 259211 | 0.997481 |
